# Towards key principles of host-associated microbiome assembly

**DOI:** 10.1101/2025.10.23.684190

**Authors:** Gui Araujo, Torsten Thomas, José M Montoya, Nicole S Webster, Miguel Lurgi

## Abstract

Symbiotic relationships between microbes and host organisms frequently involve the assembly of complex microbial communities (microbiomes) within hosts. Patterns at the community level influence life-history traits, the evolutionary trajectories of both microbes and their hosts, and are often critical for maintaining host health. These community-level patterns are dynamically driven by eco-evolutionary mechanisms acting at the individual level, such as microbial dispersal, host selection, and microbe-resource interactions. Critically, we still lack a clear picture of the ways in which these mechanisms interact to shape microbiome assembly. Here, we present a model of assembly to describe how distinct community structures can be characterised by underlying mechanisms. To illustrate the approach, we analyse microbiome data from marine sponges and simulate different structures of host-microbe associations, thereby bridging mechanistic models and empirical patterns. We further apply the model to human microbiome data to explore its relevance across biological systems, proposing that the combined effect of a small set of general mechanisms may govern diverse patterns of microbial diversity and abundance. Our findings advance ecological theory by linking individual-level processes to community-scale patterns, illuminating the key drivers underlying microbiome assembly.

---

Virtually all multicellular organisms on Earth live in close relationships with microbial communities, both influencing and being influenced by them. In many species, including humans, these microbiomes are essential for their health and survival [1]. Interactions among hosts, microbes, and their environment occur across spatial and temporal scales, thus shaping the eco-evolutionary dynamics of host-microbiome systems [2,3] and the resulting patterns of coevolution observed in these systems [4–6]. This interplay between ecological and evolutionary processes thus generates systematic differences in microbiome composition among hosts occupying distinct environments. As a result, host-associated microbial communities exhibit diverse structures and corresponding functional roles, shaped by their evolutionary and ecological contexts [7–9].

Microbiomes are typically large and diverse microbial communities that assemble throughout a host’s ontogenetic development [6]. Well-documented examples of joint host-microbiome development include the human gut microbiome [10], plant rhizospheres [11], and sponges [12]. A microbiome’s composition is shaped by a variety of biotic (e.g., ecological interactions) and abiotic (e.g., environmental properties) factors, often converging into a characteristic species-specific structure towards the end of the individual’s developmental phase [10,12,13]. These factors, or assembly mechanisms, can be quantified and incorporated into theoretical models of microbiome development that allow researchers to investigate their combined effects in shaping patterns of diversity and composition in these microbial assemblages [14,15]. Assembly mechanisms include a broad range of processes that may be mediated by the biology and ecology of microbes, host traits, environmental conditions or spatial dynamics as well as by synergies amongst them [14].

For example, the dispersal of microbes across the host-environment interface or by horizontal transmission between hosts [16,17] drives the acquisition of new microbial taxa by the host. Once transmitted, microbial interactions with the host and with other microbes can critically influence their ability to colonise and persist within the microbiome [7,18,19]. Microbial colonisation is also governed by selective pressures imposed by the host’s internal environment. These pressures can favour the enrichment of specific microbial taxa through host filtering [18] or immune selection [20], especially those beneficial to the host, thus shaping the functionality and final composition of the microbiome [7,18,21,22]. Interactions among microbes, and between microbes and hosts, are often mediated by resource exchanges shaped by microbial metabolic traits. These include cross-feeding among microbes and the bidirectional exchange of resources between microbes and their hosts [23–26]. Therefore, microbial dispersal, host selection and enrichment, and resource-based interactions represent key dynamical drivers of microbiome assembly and can thus strongly influence community-level patterns of microbiome organisation. Quantitative changes in these mechanisms have the potential to drive distinct types of communities.

Marine sponges, and the complex microbiomes they harbour, exemplify how variation in assembly mechanisms can give rise to distinct types of microbial communities across host species [8,27]. The ecological and evolutionary processes driving the assembly of marine sponge microbiomes have given rise to two broad types, that can be classified based on the total cell abundance of their associated microbiomes [28]. Low microbial abundance (LMA) sponges host relatively sparse microbiomes, whereas high microbial abundance (HMA) sponges host communities that are over a hundred times denser [28,29]. Beyond total microbial load, HMA- and LMA-associated microbiomes differ in other community-level properties, including higher microbial species richness in HMA sponges and marked differences in beta-diversity between the two groups [8,30,31]. At the host level, HMA sponges exhibit lower pumping rates [32–34], allocate more resources to their microbial communities [33], form tighter functional associations with their symbionts, and more actively enrich specific microbial taxa than LMA sponges [9,35,36]. These physiological traits (pumping rate, feeding behaviour, and microbial enrichment) enable general mechanisms of microbial dispersal and selection via resource-mediated processes, thereby shaping microbiome assembly and organisation. Thus, evolved host traits for water pumping and microbial selection, likely shaped by coevolution with their microbiomes, could help explain the divergent patterns of abundance and diversity observed between LMA and HMA sponge microbiomes.

Despite the extensive knowledge on the potential mechanisms behind the structure of different host-associated microbiomes, we still lack a unified understanding of how these mechanisms, and their interplay, give rise to patterns of organisation observed in these complex ecosystems. To this end, theoretical models of community assembly that incorporate these mechanisms can help us bridge the gap between empirical identification of individual processes and system-level understanding. Although a range of modelling approaches have recently been developed linking ecological processes to patterns of biodiversity in microbial communities [37–39], they focus almost exclusively on interactions between microbes [but see 40-42], with relatively little attention given to specific mechanisms of host-microbe interactions. Crucially, the incorporation of host-centric mechanisms is important when aiming to understand the emergence of distinct microbiomes across host or site types, as is the case in HMA vs LMA sponges, or across body sites such as gut, mouth, and palms in the human microbiome. Identifying a minimal set of assembly mechanisms that operate across biological systems, while imposing system-specific constraints through host-microbe interactions, would allow us to both unify and mechanistically characterise the emergence of distinct microbiome community structures.

In this paper, we contribute to filling this gap by developing a mechanistic theory for microbiome assembly that incorporates three fundamental processes inferred across host-microbiome systems: microbial dispersal, host selection, and microbe-resource interactions. We use this theory to investigate large-scale patterns of diversity and abundance characteristic of microbiomes, thus providing a synthetic view of the emergence of organisation in host-associated microbial communities. By incorporating empirically observed differences in pumping, enrichment, and microbial feeding across marine sponge types, our model recovers the differences in microbiome structure, including variation in richness and beta diversity, observed between HMA and LMA sponges. Furthermore, we show that a small quantitative modification to the theory (i.e. a small increase in microbial dispersal) allows us to adjust the system-specific characterisation and recover biodiversity patterns observed in data from the human microbiome across different human body sites.

## Results

Our model of microbiome assembly considers multiple host individuals, each connected to a shared external environment composed of a fixed pool of *S* microbial and *R* resource types (Fig 1A). Hosts acquire both microbial and resource types from the environment and thus assemble their individual microbiomes. Microbes consume and produce resources following a bipartite network of microbe-resource interactions (i.e. microbes do not interact directly, see Methods for details on the network structure). The model incorporates three key mechanisms that drive microbiome assembly (Fig 1B): (1) microbial dispersal, which governs the acquisition of microbes from the environment; (2) host selection, which promotes the differential growth of microbes within hosts based on their resource production; and (3) cross-feeding interactions between microbial types, in which microbes consume and release resources while relying on the availability of specific resources within the host, resulting in complex interdependencies. The model generates microbiome-level properties such as total microbial abundance, species richness (i.e. number of distinct microbial types), and species abundance distributions, allowing us to assess how the interplay among assembly mechanisms gives rise to distinct communities (Fig 1B). To simulate community assembly, we formulate a system of differential equations that tracks changes in the abundance of each microbial and resource type within each host (Fig 1C). These equations explicitly represent the three mechanisms alongside terms for microbial death, resource loss, and intraspecific microbial competition (Box 1). As an example of pattern emergence, differences in the relative strength of individual mechanisms, such as increased dispersal or stronger host selection, can lead to substantial variation in total microbial density (microbial count per volume), or abundance, across hosts (Fig 1C). These outcomes allow us to classify hosts and their assembled microbiomes into distinct community types based on emergent structural features (Fig 1D).

**Figure 1.**
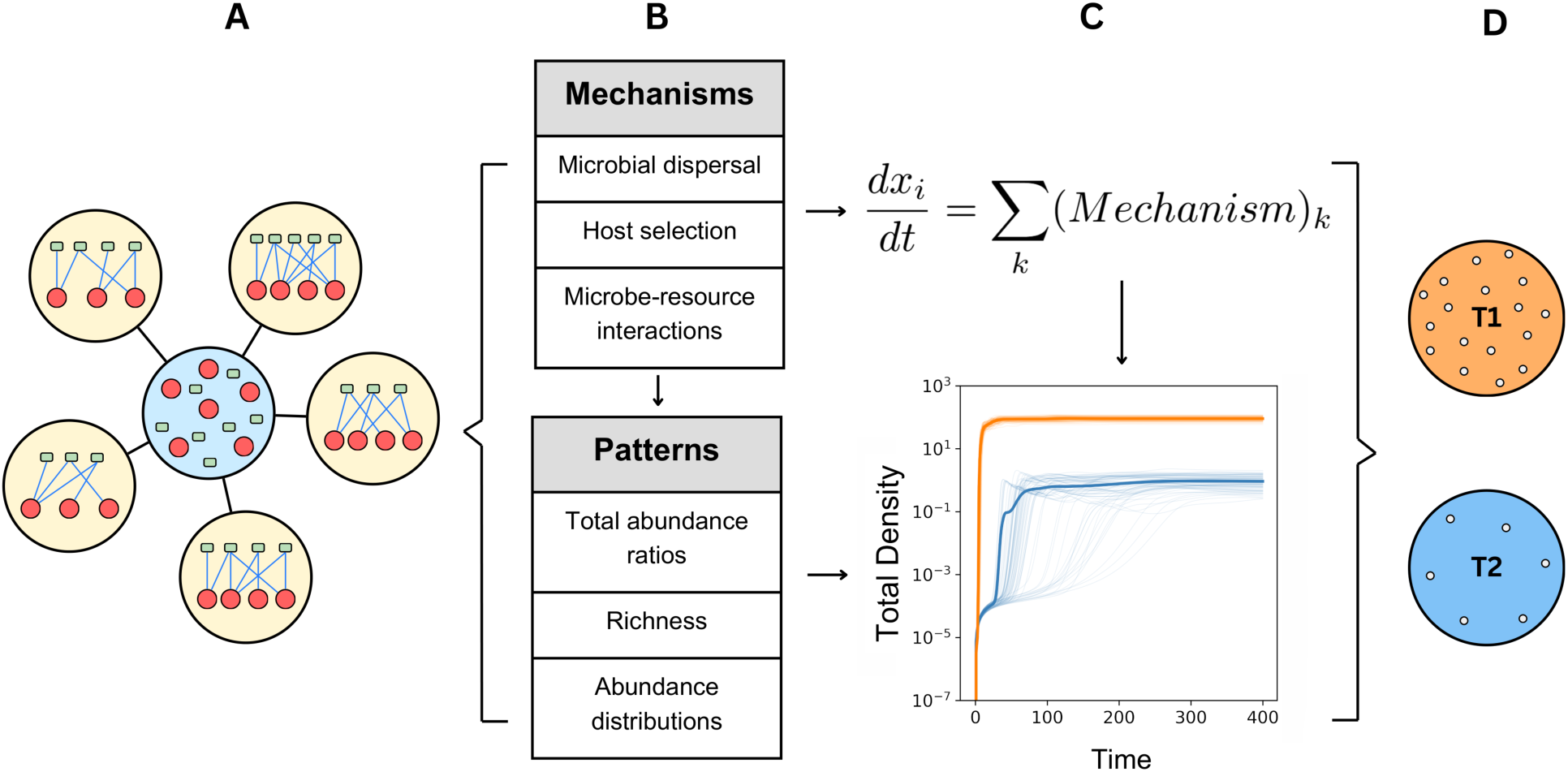
A mechanistic theory for host microbiome assembly. (A) Model setting: multiple host individuals (large yellow circles) are connected to a shared environment (large blue circle) composed of a constant pool of microbial (red circle) and resource (green square) types. Each host acquires microbes and resources from the environment and assembles a microbiome represented as a bipartite network linking microbes to the resources they consume and produce. (B) Assembly mechanisms are implemented at the host individual level and drive community-level patterns of host-associated microbiomes. Dispersal governs microbial acquisition, host selection shapes differential microbial growth, and resource-mediated interactions structure microbe-resource network dynamics. (C) The microbiome assembly is modelled through a system of differential equations incorporating various mechanisms and describing temporal changes in microbial and resource densities (microbial counts per volume), or abundances, within each host. Differences in the strength of the assembly mechanisms across hosts generate variation in microbiome community-level properties, such as species richness and total abundances. Each coloured trajectory represents the assembly of a microbiome within a single host, and the averages are highlighted. Different colours represent different relative strengths of the established mechanisms. (D) As a result, hosts can be classified into distinct community types, such as T1 and T2 for high and low total microbial abundance, respectively.

### Box 1. Ecological model of microbiome assembly

Our theory considers individual hosts that import microbial and resource types from a shared environment at a certain rate. Hosts selectively promote the growth of microbes that match their metabolic requirements. Each microbial type consumes a specific set of resources. A fraction of these resources is used for reproduction and the rest is converted into another set of resources interpreted as metabolic byproducts. Let *x*_*ih*_ denote the abundance of microbe *i* in host *h*, and *r*_*lh*_ the abundance of resource *l* in host *h*. The ecological dynamics within each host are governed by the following system of differential equations:

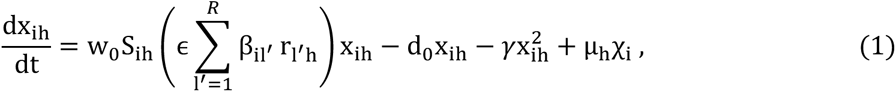

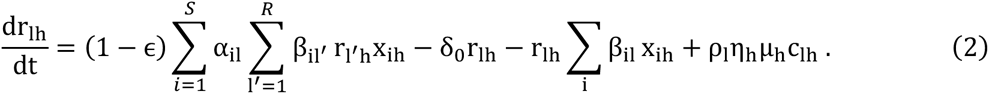

where primes denote dummy variables under summation (to avoid confusion with the focal resource *l*) and each term has a defined ecological interpretation:

#### 1. Microbial reproduction

*w*_0_*S*_*ih*_(*ε* ∑_*l*_’*β*_*il*’_ *r*_*l*’*h*_)*x*_*ih*_: Microbial growth depends on an intrinsic reproduction rate *w*_0_, the strength of host selection *S*_*ih*_, and the availability of consumed resources. Consumer-resource interactions are determined by consumption rates *β*_*il*_’ of microbe *i* on resource *l*, and the effect on growth is summed over all consumed resources. Conversion efficiency of consumed resources into new biomass is given by *ε*.

#### 2. Microbial loss and competition

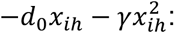 Microbes experience constant mortality at rate *d*_0_, and intraspecific competition at rate *γ*, which applies uniformly across microbial types. Interspecific competition is mediated indirectly via shared resources.

#### 3. Microbial dispersal from the environment

*μ*_*h*_*χ*_*i*_: Microbes from a well-mixed external pool enter the host at a rate *μ*_*h*_, where *χ*_*i*_ is the constant environmental abundance of microbe *i*.

#### 4. Production of metabolic by-products

(1 − *ε*) ∑_*i*_*α*_*il*_ ∑_*l*’_ *β*_*il*’_ *r*_*l*’*h*_*x*_*ih*_: The fraction of consumed resources not used for reproduction is transformed into metabolic products. The rate at which microbe *i* produces a specific resource *l* from the total being produced is given by *α*_*il*_, which must satisfy ∑_*l*_ α_*il*_ = 1. If *l* is not produced by microbe *i*, then *α*_*il*_ = 0.

#### 5. Resource depletion

−*δ*_0_*r*_*lh*_ − *r*_*lh*_ ∑_*i*_ *β*_*il*_*x*_*ih*_: Resources degrade at a constant rate *δ*_0_ and are depleted by microbial consumption, with uptake rates defined by *β*_*il*_.

#### 6. Resource uptake from the environment

*ρ*_*l*_*η*_*h*_*μ*_*h*_*c*_*lh*_: Hosts absorb resources from the external environment, where *ρ*_*l*_ is the environmental concentration of resource *l*, *μ*_*h*_ is the host’s uptake rate, *c*_*lh*_ the uptake efficiency per resource, and *η*_*h*_ the fraction of absorbed resources allocated to the microbiome.

In our simulations, we assumed three key differences between groups of hosts composed of slightly varying individuals. The first is the microbe and resource uptake rate µ_*h*_. The second is the strength of microbial enrichment *S*_*ih*_ through distinct selection capacity *v*_*h*_ (Box 2), and the third is the allocation of resources to the microbiome, represented by η_*h*_ as the fraction of acquired resources that is available to microbes within the host. The set of *q*_2_ non-essential resources (Box 2) characterises the distinction at the host species level. Relatively smaller individual differences across individuals within host species were modelled as small random variations in trait values (see Methods).

### Box 2. Microbial enrichment within hosts

Host individuals require resources that are not available in the environment and can only be synthesised by microbes (i.e., secondary and tertiary metabolic products, see Methods for details on the microbe-resource network and classification). We assume that all hosts require a shared set of *q*_1_ randomly defined essential resources drawn from secondary and tertiary resources, which are valued by all host species (universally valued). In addition, each host species is characterised by valuing a particular set of non-essential resources, drawn randomly from the remaining secondary and tertiary resource pool. Each host has an equal probability of valuing any given non-essential resource, resulting in an average of *q*_2_ additional valued resources per host species. This variation in host requirement shapes species-specific selection pressures on microbial community composition.

Each host has a selection capacity *v*_*h*_, defined as the probability of selecting any microbial type that produces at least one resource valued by the host. When a microbial type is selected, its reproductive advantage increases with the number of valued resources it produces. For example, a microbe producing three valued resources is more beneficial, and therefore more strongly enriched, than one producing only a single valued resource. This defines a quantitative, function-based selection mechanism.

To model the greater adaptive value of microbial specialisation, we assume a superlinear selection curve, where the strength of selection increases faster than linearly with the number of valued resources. If microbe *i* produces *t*_*ih*_ resources valued by host *h*, and *θ*_*ih*_ is a binary variable indicating whether the microbe has been selected (*θ*_*ih*_ = 1 with probability *v*_*h*_ when *i* produces at least one resource valued by *h*, otherwise 0), then the selection factor *S*_*ih*_ affecting microbe *i*’s reproduction in host *h* is given by:

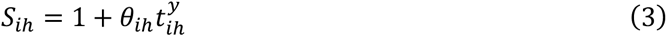

When a microbe is not preferred by the host (*θ*_*ih*_ = 0) or does not produce any valued resources 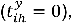 then *S*_*ih*_ = 1, meaning no enrichment. *y* is a parameter that controls the rate of increase in the response of *S*_*ih*_ to the production of valued resources and is set to *y* = 2 in our simulations to implement moderate superlinearity.

### Microbial dispersal, enrichment, and resource dynamics drive the assembly of distinct host types

Using the theory described above, we set up to investigate the ability of mechanisms identified by previous studies as potential drivers of sponge microbiome structure in recovering observed organisational features of these communities. We thus parameterised our model based on known biological features of marine sponges. Marine sponges are able to control the import of microbes and resources from the external environment via pumping of seawater, thereby driving microbial dispersal [43]. Empirical data have shown that LMA sponges exhibit ∼2.5 times higher pumping rates (µ_*h*_ in our model) than HMA sponges [33]. We thus incorporate this heterogeneity across host types into our model. In terms of the host selection mechanisms, it is known that some sponges (e.g., HMA sponges) can maintain more intimate and functionally specific associations with their microbiomes than others [9,35,36]. We modelled this phenomenon by allowing differences in the selection capacity of microbes by hosts (*v*_*h*_), resulting in a greater probability of enriching microbes that produce valuable metabolic resources in hosts with higher *v*_*h*_ values. Finally, substantial differences in the magnitude of resource allocation have been observed across sponge types. HMA sponges typically allocate a greater share of environmental resources to their microbes, consuming less resources directly. To reflect this, in our model, we established 100-fold differences in the fraction of resources made available to microbes (η_*h*_) between groups of hosts, in line with empirical estimates for differences between HMA and LMA sponges [33].

We simulated communities comprising multiple host species of both LMA and HMA types (see Methods for full specification of parameter values), with 72 individuals in each group, divided into 12 species of 6 individuals each, and implementing the distinctions described above across host types for all three mechanisms. We contrasted our model results with empirical patterns of sponge-associated microbial communities where sponge samples from many species had previously been classified into HMA and LMA categories based on total microbial biomass in host tissues [44]. To make our simulated data comparable to empirical data, we sampled microbial counts proportional to final abundances (24,000 counts per sample) to generate simulated sample x species tables (equivalent to the Operational Taxonomic Units (OTU) tables commonly analysed in microbiome studies) [27].

Our model produced a difference of two orders of magnitude in total microbial abundances between HMA and LMA samples, consistent with empirical findings reporting microbiomes at least 100 times more abundant in HMA sponges [28,29] (Fig. 2A). Composition of simulated communities also formed two distinct clusters corresponding to HMA- and LMA-type hosts, as revealed by non-metric multidimensional scaling (NMDS) on Bray-Curtis dissimilarities (Fig. 2B) and PERMANOVA tests confirming that variation between groups was significantly greater than variation within groups (pseudo-F = 18.5, permutation p = 0.001).

**Figure 2.**
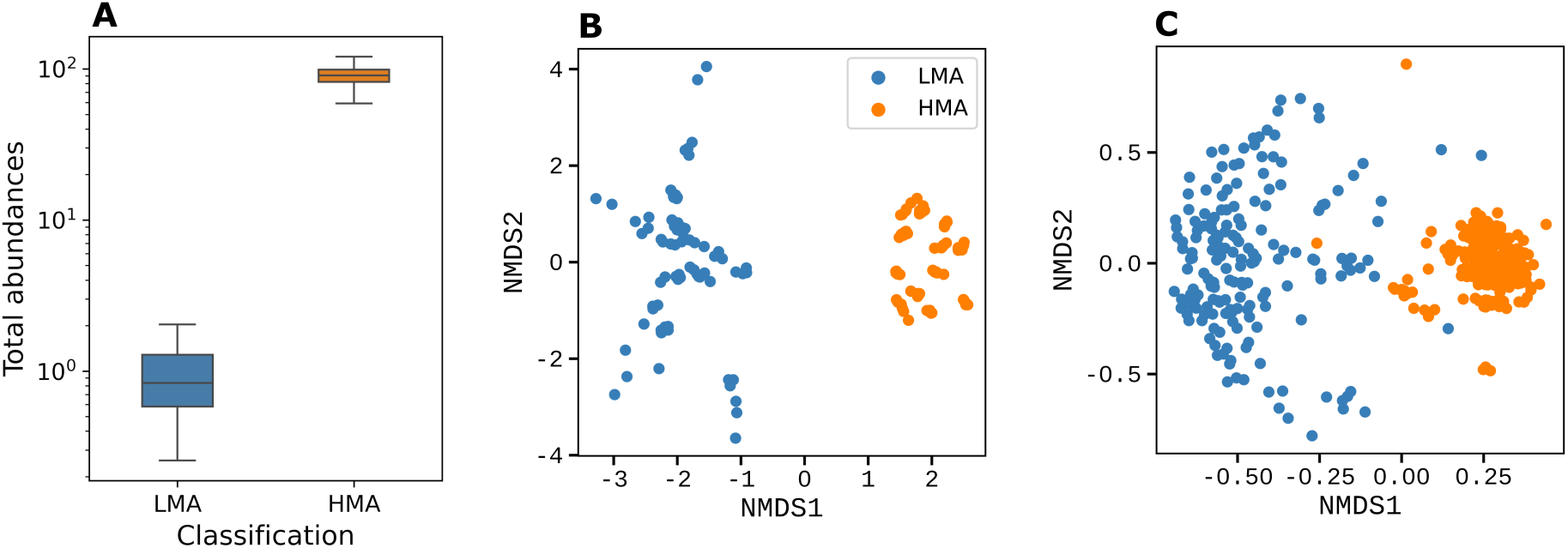
Compositional differences observed between microbiomes of HMA and LMA sponge hosts are determined by microbial dispersal, host selection and enrichment. Comparison of model outputs and empirical data (Operational Taxonomic Unit or OTU tables) of marine sponge samples from the Sponge Microbiome Project (see Methods for details): total abundances (only simulation) and community composition of microbiomes highlighting the separation between groups of host types. (A) Simulations of several LMA- and HMA-type hosts after microbiome assembly reproduce the defining characteristic of the HMA-LMA separation. Samples distributions from the model show a difference of two orders of magnitude in the total abundance (or density) of microbes within simulated host individuals separated by their classification (72 samples of each). (B) Non-metric multidimensional scaling (NMDS) of Bray-Curtis dissimilarities in relative abundances of OTU-equivalent units in the simulated communities. OTU-like tables were generated by sampling modelled microbial types with probabilities proportional to their final abundances. The analysis reveals two well-separated clusters corresponding to HMA- and LMA-type hosts. (C) NMDS applied to empirical data also shows a clear separation between HMA and LMA samples. Rarefaction of the empirical data was set at 23,376 counts (minimum number of reads in the dataset) on 312 HMA and 184 LMA samples divided into 19 HMA and 17 LMA species. The model was sampled at 24,000 counts, generating simulated tables for 72 HMA and 72 LMA samples, equally divided into 6 HMA and 6 LMA species. In the model, LMA-type hosts had pumping rates 2.5 times higher than HMA-type hosts, resource allocation to microbes was 100-fold lower in LMAs, and selection capacity was set to 0.01 for LMA and 0.2 for HMA hosts.

This clear separation mirrors that observed in empirical sponge microbiome data, calculated from 312 HMA and 184 LMA samples divided into 19 HMA and 17 LMA species and rarefied to 23,376 reads (Fig. 2C) (PERMANOVA tests: pseudo-F = 77.8, permutation p = 0.001). A further similarity lies in the within-group spread: LMA samples showed greater dispersion, while HMA samples were tightly clustered in both modelled and empirical data (LMA to HMA ratio of mean distance to centroid of around 1.80 in the entire data, 1.38 in a subsample of data with similar size to the model shown in Fig S1, and 1.16 in the model). An analysis of a subsample of the data with a sampling size closer to the model and more homogeneous between groups (7 HMA and 7 LMA species with 6 samples each) reveals no meaningful changes (Fig S1). These results demonstrate that the assembly model successfully reproduces the defining features of the observed dichotomy between HMA and LMA sponge microbiomes, lending support to the hypothesis that the identified mechanisms are likely drivers of assembly in host-associated microbiomes.

Each of the mechanisms above were tested in isolation to determine their relative contributions to observed empirical patterns. Interestingly, none of the mechanisms in isolation was able to reproduce empirical patterns of microbiome organisation. For instance, we tested differences in pumping rates as the only distinction between host groups, with parameters concerning the other mechanisms remaining equal. This resulted in lower abundance and richness for the HMA-type hosts (i.e. those with lower pumping rate), contrasting sharply with empirical observations (Fig S2A). Higher resource allocation and selection capacity would then be required to counterbalance the lower pumping rate, allowing the HMA’s more stable internal environments and lesser variation across microbiomes [8,9]. We tested each of these two mechanisms in isolation counteracting lower pumping rates, which resulted in partial matches to empirical observations (Fig S2B and S2C). Therefore, the action of both mechanisms together, on top of a lower pumping rate, were necessary to reproduce the patterns seen in natural microbiomes. These tests suggest that hosts with lower pumping rates can afford to absorb less microbes and resources from the environment if they are better able to exert a tighter control on their internal environment, in a trade-off that allows them to draw more benefits from their microbiome.

### Beyond compositional differences: species richness and abundance distributions

Previous analyses of marine sponge microbiomes have revealed that HMA sponges typically harbour more complex microbiomes, both in terms of the absolute number of species (or OTUs) per host as well as the evenness of their abundances, than LMA sponges [31] (Fig 3A,C), a distinction that is well captured by our model (Fig 3B,D). Further, rank-abundance curves from empirical data reveal greater variation in abundance distributions among LMA sponges (i.e. size of shaded area around the lines in Fig 3E; [31]). Moreover, HMA microbiomes tend to show slightly greater similarity in the abundances of dominant taxa, with a smaller slope of the curve for higher abundances suggesting a higher degree of shared dominance of abundant microbial types. These trends are also recovered by the model, with the HMA curve being more horizontal than the LMA curve at the beginning (Fig 3F). Altogether, these results indicate that differences in microbial dispersal, host selection, and resource allocation implemented in the assembly model can jointly reproduce observed patterns of richness, evenness, and abundance distributions in HMA vs LMA microbiomes.

**Figure 3.**
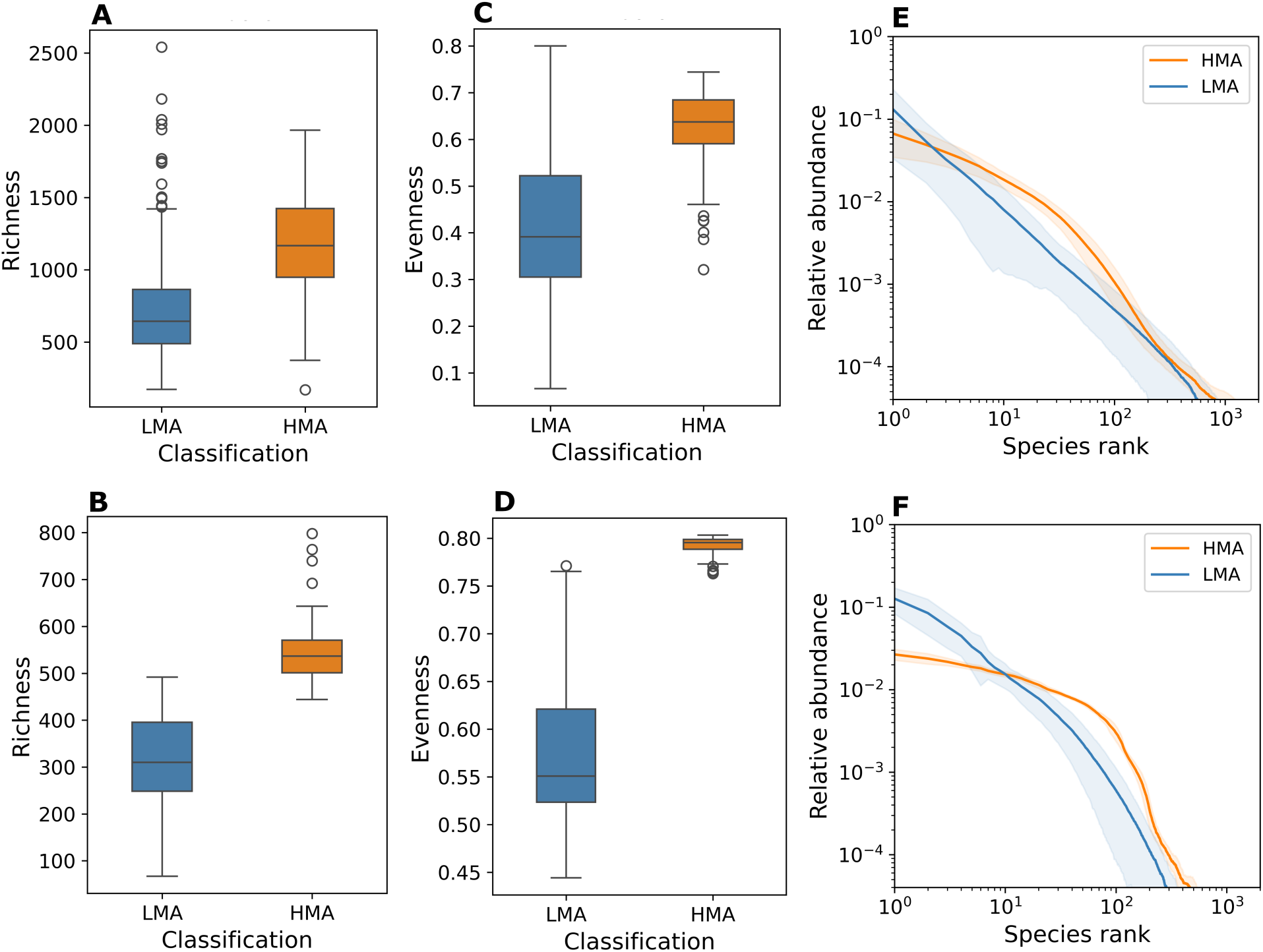
Microbiome complexity is driven by differences in microbial dispersal, selection and enrichment across host types. Comparison of microbiome complexity between empirical data (OTU tables) of marine sponge samples from the Sponge Microbiome Project (top row) and model simulations (bottom row). Box plots show the distribution of host samples generated from the OTU tables, empirical (312 HMA and 184 LMA samples) and simulated (72 HMA and 72 LMA samples), separated by their classification. (A,B) HMA sponges exhibit higher number of OTUs or microbial species richness than LMA sponges in natural samples. The same relationship is preserved in simulations. While the total pool of OTUs in the model is S = 3,000, the natural data includes 56,161 OTUs. (C,D) Evenness is lower and more variable among LMA sponges in the empirical data. Model outputs reproduce this pattern, with HMA samples exhibiting consistently higher evenness. (E,F) Rank-abundance plots of average relative abundances (i.e. the sample averages of each rank) show differences in abundance distributions across host types. HMA hosts distributions present a smaller standard deviation (shaded region, the sample standard deviation of the ranked abundances) and exhibit a more prominent curve of abundances, representing more similar distribution of taxa with higher abundances across both empirical and model data. Simulations were the same as in Fig 2.

To assess the sensitivity of the model to the functional form of the host selection function *S*_*ih*_, we ran additional simulations for several values of the exponent *y* = {0.5,1, 1.5, 2.5, 3} and obtained similar results, although the group spread in ordination analysis suggests that values around 2 best reflect the data (Fig S3). We also assessed the sensitivity of the model to the number of fixed and random resources valued by host individuals (Box 2). The main model uses (*q*_1_ = 15, *q*_2_ = 15) and we tested different proportions and values: (*q*_1_ = 15, *q*_2_ = 0), (*q*_1_ = 15, *q*_2_ = 30), (*q*_1_ = 5, *q*_2_ = 15), (*q*_1_ = 30, *q*_2_ = 15), and (*q*_1_ = 30, *q*_2_ = 30). Similar results were obtained in all cases, although the group spread in ordination analysis suggests that similar proportions better reflect the data (Fig S4).

### Unifying assembly mechanisms across microbiomes

In marine sponges, pumping and microbiome feeding constitute physiological processes through which ecological mechanisms of dispersal and resource dynamics are manifested. Other host-microbiome systems might differ in biological details but are likely to incorporate processes with similar roles in the generation of mechanisms of acquisition, selection, and microbe-resource interactions. Therefore, the dynamical model developed above should be able to capture community-level patterns observed across host-microbiome systems. To explore this, we analysed time-series data extracted from human microbiomes from different body sites, including tongue, faeces, and palm [45,46] and contrasted empirical patterns from this system with our theoretical expectations. Based on the assembly mechanisms proposed in our theory, we expected that palm microbiomes could be distinguished from tongue and faeces due to the fundamental differences of the former being external to the body, and thus more strongly subjected to environmental influences, as well as not being part of an internal body system (i.e. the digestive system). We hypothesised that, as for the LMA group, the palm microbiome experiences a higher turnover of microbial cells and smaller resource allocation and selection by the host.

We confirmed the differences in microbial species richness, evenness and abundance distributions between the palm and the tongue and faecal microbiomes (Fig 4A-C). However, contrary to the microbiomes of LMA sponges, those of palms comprise the group with higher richness and evenness when compared to the tongue/faeces group (see Fig S5 for direct comparisons between datasets). Interestingly, using the same model setup as before, and only increasing the differences in microbial dispersal (palm having 10 times the tongue/faeces dispersal instead of 2.5 as used for the LMA against HMA sponges), our model more closely approximates the observed differences across human microbiomes (Fig 4D-F). Although simulation results for evenness could be closer to observations, the model was able to capture the trends in the variability of both groups (Fig 4C,4F). These results suggest that the difference in assembly mechanisms of palm against tongue/faeces in human microbiomes and LMA against HMA in sponge microbiomes are mainly represented by a larger difference in microbial dispersal/ colonisation from the external pool. This observation could be explained by the human palm being typically exposed to a more diverse array of contributing input environments (i.e. humans touch many different surfaces on a daily basis) as compared to the relatively more constant input of seawater experienced by sponges, given their sessile life.

**Figure 4.**
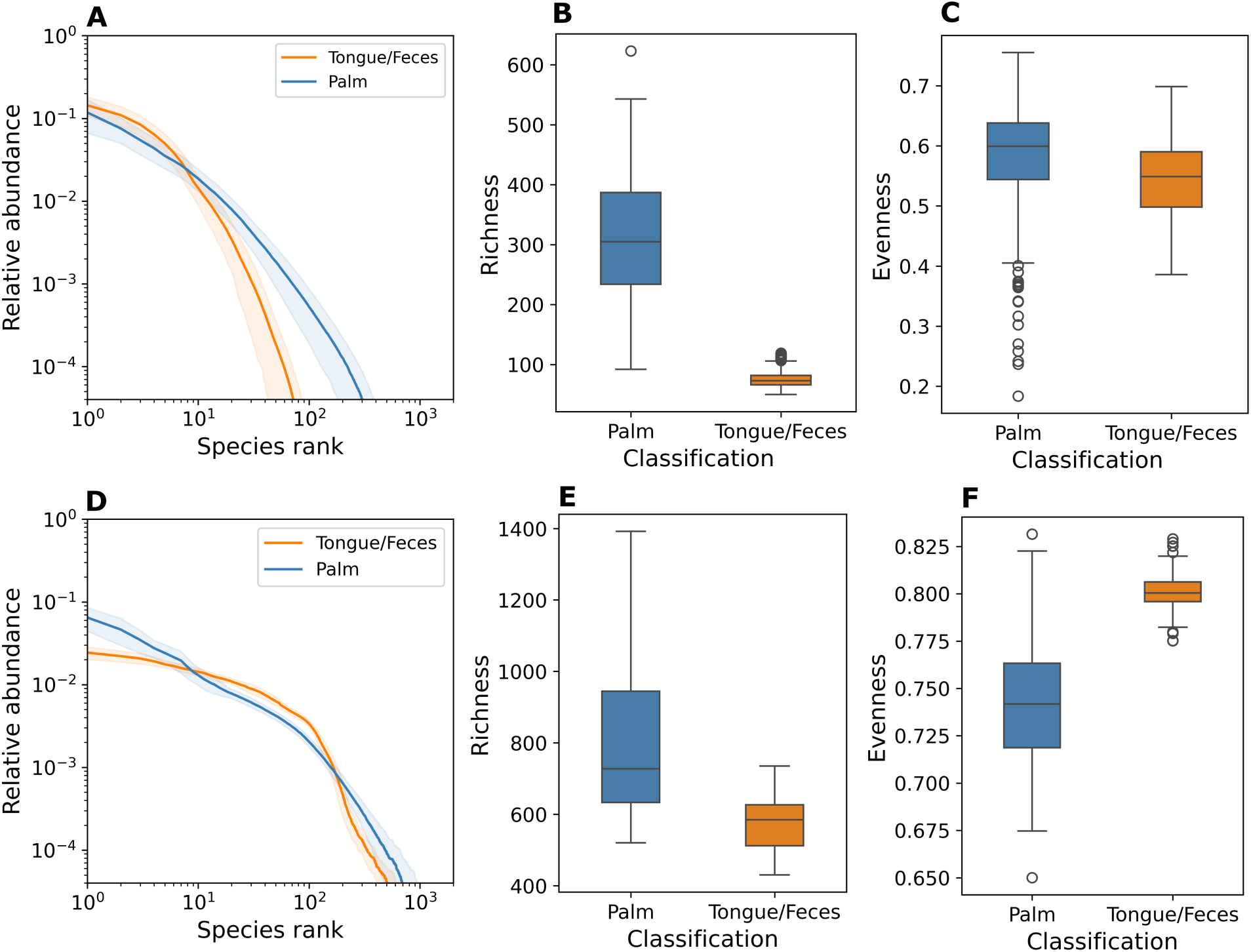
Human microbiome analysis suggests generality of assembly mechanisms across microbiome types. Comparison of different human microbiome sites (top row) with model outputs (bottom row) shows how the same assembly mechanisms, with quantitative parameter changes, can generate distinct microbiome patterns observed across systems. Empirical timeseries data of microbiomes from palm (two sets of 265 and 20 samples), tongue (319 samples), and faeces (330 samples) of a human subject available from MGnify EBI Metagenomics (see Methods for details). Simulations included 72 samples each for palm- and tongue/faeces-type communities, ran over 250 time units (sufficient for microbiome assembly in this case). Box plots show the distribution of host samples generated from the OTU tables, empirical and simulated, separated by their classification. (A) Rank-abundance curves for tongue, faeces, and two palm datasets show their mean relative abundance (solid line, see previous figure for details) and standard deviation (shaded region). Curves are coloured by group. The palm microbiome (external body surface) forms a distinct cluster from the tongue and faecal microbiomes (internal body environments, both part of the digestive system), which show similar abundance profiles. (B,C) Palm microbiomes exhibit consistently higher richness and slightly higher evenness than those from the tongue or faeces. (D-F) Simulation results approximate the empirical patterns, closely resembling observed trends in relative values and variability. To theoretically investigate empirical patterns of human microbiomes, tongue/faeces microbiomes were modelled as having a higher microbial enrichment and resource availability compared to palm microbiomes, analogous to HMA (vs LMA) sponge hosts. Palm microbiomes were given a 10-fold higher microbial flux relative to tongue/faeces (versus the 2.5-fold ratio used for LMA vs. HMA sponges). The data was rarefied with 20,000 reads, and we sampled the model to generate tables with the same number of counts. See Methods for further details.

## Discussion

Mechanistic modelling of host-microbe associations, grounded in quantitative representations of biological mechanisms observed in empirical systems, offers a powerful theory to better understand the factors governing the emergence of these complex relationships. This is a growing field which in the last decade has revealed fundamental insights into the ways in which nature can shape complex microbial associations, both free living [39,47,48] and host-associated [38,40,49]. This work has revealed underlying general mechanisms of microbial interactions that can result in connectivity patterns, and their dynamical consequences, enhancing the stability and persistence of microbial assemblages. In host-microbiome associations however, specific controlling mechanisms of the host are hypothesized to influence microbiome dynamics [9,14,33,41]. Incorporating these processes into a general theory of host-associated symbioses is fundamental to better understand their formation.

In this paper, we introduced a general ecological model and theory of community assembly that links host-microbiome interactive mechanisms to emergent patterns of microbiome organisation and diversity. The model incorporates microbial dispersal, host selection and enrichment, and resource-mediated microbial interactions as fundamental underlying mechanisms of microbiome assembly. These factors are sufficient to capture variation in total abundance, richness, and abundance distribution of taxa across microbiomes of different host types. But the processes need to operate in conjunction-each, on their own, are unable to capture empirical realism. We used this theory to investigate known differences between two marine sponge host types, high and low microbial abundance (HMA and LMA), providing an answer to the question: What ecological mechanisms give rise to the group-level separation observed across types? Moreover, we have shown that this baseline theory is broadly applicable across diverse host-microbiome systems. Relatively small modifications to quantitative values of the model parameter controlling microbial dispersal were sufficient to apply our theory to investigate differences observed across human microbiomes. In this way we were able to explain differences observed between human palm and digestive system microbiomes. Our findings suggest the existence of general underlying principles being key to the assembly of microbiomes across taxonomically diverse host organisms.

Local environmental conditions and the dispersal capacity of individuals and species have long been recognised as fundamental mechanisms driving biodiversity across ecological and spatial scales [50,51]. Our theory lends support to the accumulating evidence that these processes are indeed fundamental for the assembly of microbial communities [52]. In marine sponges, we have shown that the trade-off between reduced microbial immigration (via e.g., lower pumping rates) and enhanced control of the internal host environment are central processes to maintain the HMA sponge strategy. Moreover, our results suggest that these same mechanisms are likely behind the maintenance of the human faecal/tongue microbiome and the differences observed with the more exposed palm microbiomes. Interestingly, among the mechanisms studied, resource allocation (i.e. how many resources acquired by the host are used by associated microbes) had the strongest controlling effect on the emergence of distinct microbial communities across host types, while host selection primarily refined these outcomes by improving quantitative agreement with empirical data (see Fig S2). This suggests that the mechanism of resource allocation to the microbiome is critical to the emergence of differences between functionally important vs more open microbiomes, as is the case in the HMA-LMA dichotomy in marine sponges and the digestive system versus palm microbiomes in humans. Nonetheless, the large difference in resource allocation across microbiome types (two orders of magnitude) highlights the importance of quantifying the relative strength of different mechanisms. This point is further supported by our results on human microbiomes, since changing only the order of magnitude of the microbial influx (same as the pumping rate for sponges) can considerably affect the differences between community types.

An interesting emergent property of the HMA sponge and the human digestive system groups, observed in both empirical data and simulations, is the lower variability in abundance distributions across replicate samples (Figs 3F & 4D), which is consistent with strong environmental control. Model simulations also predict a faster microbiome assembly (i.e. shorter transient dynamics to reach equilibrium) in HMA compared to LMA sponges (Fig 1C), suggesting possible testable differences in priority effects and ontogenetic microbiome development between the two groups. Faster assembly through rapid microbial enrichment may reduce colonisation by taxa arriving in later stages [53], reinforcing internal environmental control and contributing to more predictable microbiome composition. This effect is likely intensified by the higher prevalence of vertical symbiont transmission in HMA sponges [54,55], further distinguishing their assembly process from that of LMA sponges.

Host-mediated selection of microbes is a complex process that may involve the host’s immune system, biochemical interactions, the presence of defence molecules, or the creation of specialised niches that enrich or suppress specific microbial taxa [56–58]. In our theory, we have modelled host-mediated selection through a simplified enrichment function based on microbial resource production, an abstraction that captures the outcome of host selection without explicitly modelling its biological details. Despite this simplification, our results are robust to variation in the shape of the selection function, as shown in our sensitivity analysis. Qualitative patterns of microbiome diversity are preserved across different selection functions, with exponents around two providing the best fit to empirical data (Fig S3). This suggests that enrichment of microbes involves some non-linear selection process. This hypothesis could be tested experimentally by decoupling microbial functional contribution from reproductive success and quantifying their respective effects. Future modelling efforts may explore more detailed implementations of host selection to evaluate the impact of different underlying mechanisms.

In our model, microbial dispersal was modelled exclusively as horizontal transmission: hosts acquire microbes from the external environment and start the assembly process without an initial microbiome. However, vertical transmission, i.e. the microbial transfer from parent to offspring, is a well-documented process in many systems [59,60]. It not only seeds the microbiome but often involves the preferential transfer of specific microbial taxa, resulting in differential transmission [8,61,62]. This mechanism may be essential for accurately capturing microbiome composition and functional inheritance and is particularly relevant in the context of host-microbe coevolution [8,9]. Nonetheless, our results suggest that vertical transmission is not necessary to reproduce key patterns of richness and diversity across communities, at least under the ecological scenarios considered here.

Despite the potential generality of our theory in explaining differences between microbiome types, it remains limited by its reliance on *a priori* specification of the ecological mechanisms responsible for those differences. Importantly, our model does not account for how such mechanistic distinctions emerge naturally. Divergence between host-microbiomes has likely arisen through evolutionary time via eco-evolutionary processes [42,63], thus opening a very important avenue for future research: the eco-evolutionary emergence of the mechanisms of assembly. In the case of marine sponges, for example, it has been hypothesised that HMA sponge lineages evolved multiple times independently from LMA sponge ancestors across diverse clades, representing a more recent evolutionary strategy [9]. Future research could explore evolutionary scenarios in which hosts and microbes coevolve, leading to the spontaneous emergence of distinct microbiome types. For sponges, this would involve simulating the gradual divergence of host traits over generations to identify conditions under which HMA-like microbiomes arise as a discrete class rather than along a continuous gradient of coexistence. Such a study could resolve the observed group-level separation in microbiome diversity in more mechanistic detail, incorporating host phylogenetic branching and potentially linking it to genetic distances among species [64].

Our study demonstrates that mathematical models of community assembly can shed light on the drivers of emergence of complex microbiomes by accounting for differences in host type-level ecological mechanisms. Specifically, we show that variation in microbial community types, such as those observed in HMA and LMA sponges and in palm versus digestive system human microbiomes, can be explained by a relatively small number of mechanistic drivers. These findings advance ecological theory by providing a mechanistic link between individual-level processes and community-level patterns in microbial communities, bridging the gap between host traits and microbial diversity. Our work underscores the central roles of microbial dispersal, host selection, and microbe-resource interactions in shaping host-associated microbiomes. Furthermore, our results provide an improved understanding of the relative importance of different processes for the maintenance of complex microbiomes, thus allowing for the further development of theoretical approaches to effectively manipulate microbiomes, which can have practical applications for the preservation of host health and ecological communities [65,66].

## Methods

### Microbe-resource network

To capture the flow of resources through a simplified metabolic cascade, we imposed a minimal three-layer trophic structure on the microbial community that can represent cross-feeding relations respecting a progressive depletion of nutrient values. We divided microbial types into three functional groups, each corresponding to a different trophic position in the consumer-resource network:

1. The first group consists of microbes that consume primary resources (i.e. those directly imported from the environment by the host) and convert them into metabolic products.
2. The second group consumes these products (secondary resources) and produces additional compounds (tertiary resources).
3. The third group consumes tertiary resources but does not produce further usable products.

This layered organisation also defines three corresponding sets of resources: primary (externally acquired), secondary (first metabolic products), and tertiary (second-level products). As a result, the consumption (β) and production (α) matrices used in the model are structured into three distinct blocks, each representing a trophic group.

### Simulations

Each microbial type is defined by the specific set of resources it consumes and produces, represented by the matrices β_*il*_ and α_*il*_, respectively. Two microbes belong to the same type if they consume and produce identical sets of resources. To focus on host-mediated mechanisms, we standardised microbial traits across types. All microbes share the same intrinsic reproduction rate (*w*_0_), death rate (*d*_0_), and intraspecific competition coefficient (γ).

We assume that each microbial type consumes *n*_*c*_ = 100 resources and produces *n*_*p*_ = 100 metabolic products. Resource production is uniformly distributed across the selected products, with no preference for particular resource types. Thus, nonzero entries of the production matrix α take the value 1/*n*_*p*_.

To produce the results presented on Fig 2 and Fig 3, we simulated 24 sponge species divided into 12 HMA and 12 LMA species. Each species was composed of 6 individuals. We set the number of resources *R* = 5000 and the number of microbes *S* = 3000. Resources and microbes were equally separated into the three trophic groups.

Parameter values for the ecological equations were defined as follows: *w*_0_ = 5, ɛ = 0.7, *d*_0_ = 0.5, γ = 1, δ_0_ = 0.1. β_*il*_ ∼ *U*[0,10] for consumed resources and zero otherwise, ρ_*l*_ ∼ 30 ∗ *Lognormal*(0,1), and χ_i_ = 0.0001*U*_*h*_[0,1]. *c*_*lh*_ = *U*_*l*_[0,1]h1 + *N*_*h*_(0,0.1)j for primary resources and zero for the rest (values are the same for individuals of the same species). Pumping rates are µ_*h*_ = 0.0002(1 + 0.5*U*_*h*_[−0.5,0.5]) for HMA sponge samples and 2.5 times larger for LMA sponge samples. Resource allocation is η_*h*_ = 0.5(1 + 0.5*U*_h_[−0.5,0.5]) for HMA sponge samples and η_*h*_ = 0.005(1 + 0.5*U*_*h*_[−0.5,0.5]) for LMA samples. Selection capacity is *v*_*h*_ = 0.2 for HMA sponges and *v*_*h*_ = 0.01 for LMA sponges. Every sponge species values the same set of *q*_1_ = 15 essential resources and each species values on average *q*_2_ = 15 non-essential resources (the chance of valuing any additional resources is *q*_2_/ ξ, where ξ = (2*R*/3) − *q*_1_ is the number of non-essential secondary/tertiary resources).

All hosts start without microorganisms, so initial abundances are *x*^0^ = 0, and the same is true for resources. We integrate the model for 400 periods, each with 10 timesteps of size 0.01. Then, we analyse the model’s final state.

To produce the results in Fig 4, we simulated 24 sample types (analogous to sponge species) divided into 12 palm and 12 tongue/faeces types. Each type was composed of 6 individuals. The only difference from previous simulations it that microbial dispersal (same as pumping rates, µ_*h*_) of palms was 10 times that of tongue/faeces.

### Sponge microbiome data

We used the microbiome data of operational taxonomic units (OTU) from the Sponge Microbiome Project [44]. Moitinho-Silva et al. (2017) [31] used this data and presented a table of sponge sample IDs and their classification as HMA or LMA sponges based on an electron microscopy analyses [29], which we used to select sponge species and their OTU composition. We also removed rare OTUs that appeared at less than 10 counts over the entire OTU table. The classified OTU table comprises 496 samples (312 HMA sponges and 184 LMA sponges) separated into 36 species (19 HMA sponges and 17 LMA sponges). We rarefied the OTU data by randomly sampling to a minimum depth of 23,376 (minimum number of reads in the dataset) among the host samples. The pool of OTUs in the final data is of 56,161.

### Human microbiome data

We used data available from EBI Metagenomics: longitudinal human microbiome data (faeces, left/right palm, tongue; subject M3; Study MGYS00002184, ERP021896) from the EBI MGnify platform (project ERP021896) [45,46]. We considered samples having at least 20,000 reads, then applied rarefaction to calculate their relative abundances with 20,000 reads. We obtained a tongue dataset of 319 samples, a faeces dataset of 330 samples, and two palm datasets of 265 and 20 samples.

### Analysis

To prepare the model output for analysis, we used the final microbial abundances by the model to generate relative probabilities. Then, we applied a rarefaction with depth of 24,000 counts and created OTU tables equivalent to the SMP data for direct comparison (we used 20,000 counts for the human microbiome analysis).

We created a Bray-Curtis distance matrix and used a nonmetric multidimensional scaling (NMDS) analysis for dimensionality reduction and visualisation of distances of log-transformed relative microbial abundances. For that, we used the functions *decostand*, *log1p*, and *metaMDS* from the R package *Vegan* [67]. To compare the spread in the NMDS projection between groups, we calculated the mean distance of points to the group’s centroid (average position) for each group and analysed their ratio.

For additional evidence of group separation, we performed a PERMANOVA statistical test on Bray-Curtis dissimilarity matrices of HMA and LMA sponge samples. For that, we used the functions *vegdist* and *adonis2* from the R package *Vegan* [67].

As a measure of diversity, we calculated the evenness of the OTU distributions of each sample, which measures how far the Shannon entropy of the distribution of relative abundances is from its maximum. Therefore, evenness was calculated as the entropy divided by its maximum value, which is itself the logarithm of species richness. Therefore, 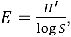 with 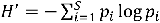 for the relative abundances *p*_*i*_. Richness was measured as the number of non-zero counts of OTUs in each sample.

For a comprehensive visualisation of the shape of abundance distributions, we ranked OTUs and plotted their ranked abundances, from maximum to minimum. We show relative abundances in the range of [10^−4^, 1]. The curves show average values with shaded regions representing the standard deviations for each rank.

## Acknowledgements

This work was supported by the Leverhulme Trust through Research Project Grant # RPG-2022-114 to ML.

## Data availability

All datasets and codes used in this study are available in the accompanying Zenodo repository https://doi.org/10.5281/zenodo.17107189.

## Supplementary figures

**Supplementary figure 1.**
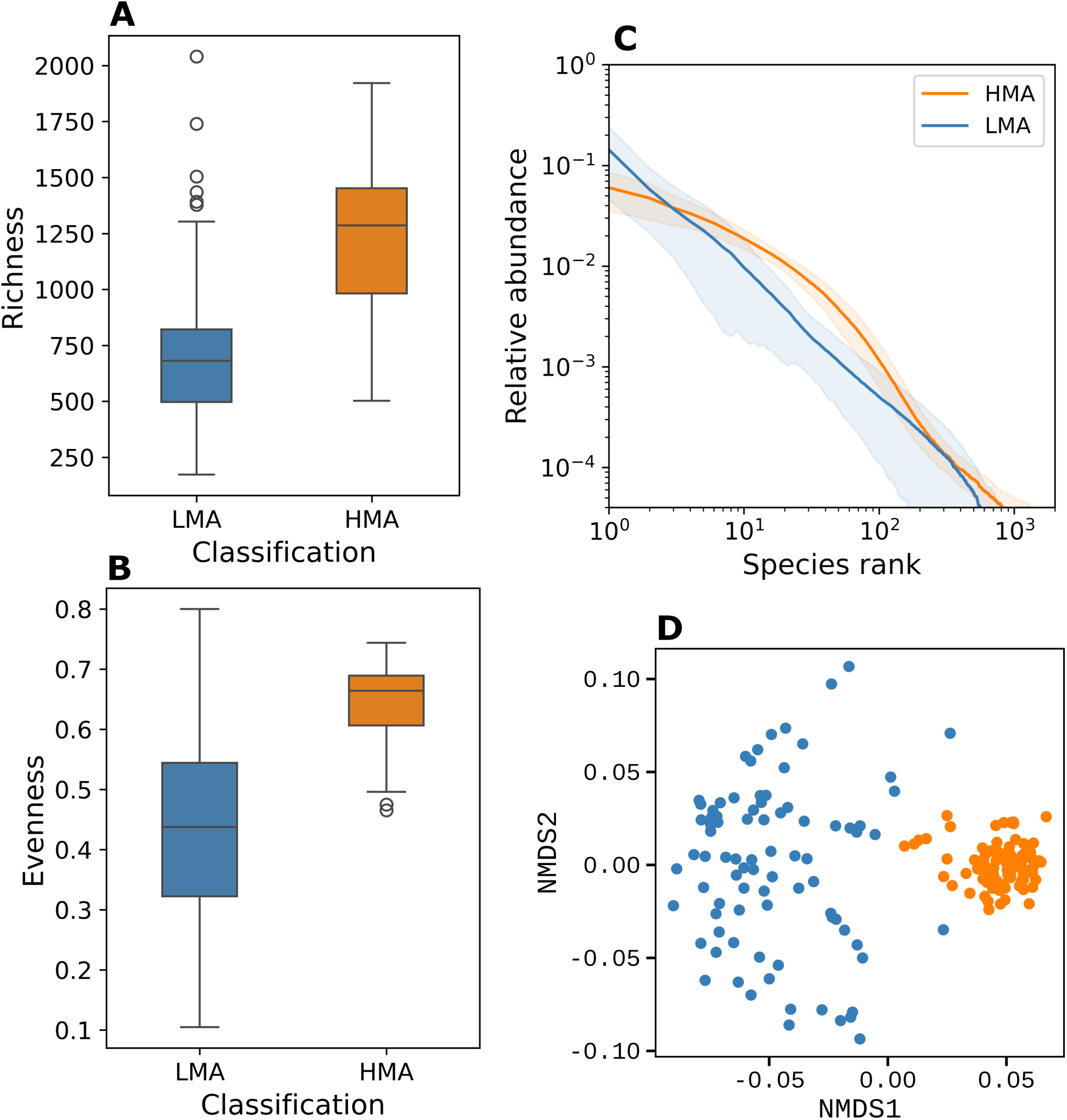
Empirical patterns of marine sponges in a subsample of the data. A subsample of the marine sponge data presented in Figs 2 and 3 to more closely match the number of samples used in simulations and to have a homogeneous sampling between groups: 7 HMA and 7 LMA species with 6 samples each. (A) Microbial richness distributions, (B) evenness distributions, (C) rank-abundance plot, and (D) NMDS ordination showing the group separation. See captions of Figs 2 and 3 for details.

**Supplementary figure 2.**
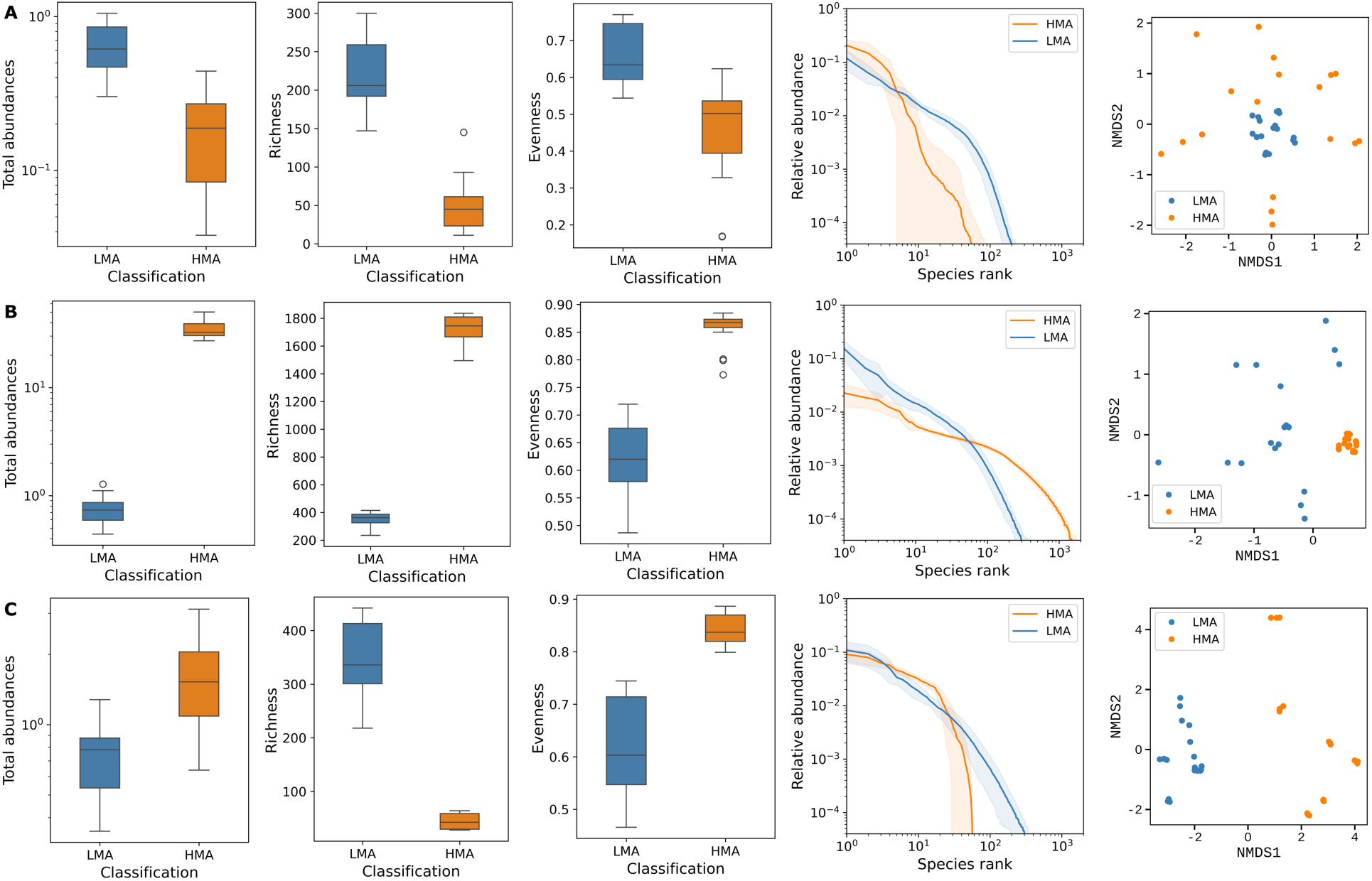
Separate effects of mechanisms. Model parameters and plots are the same as presented in Fig 2 and Fig 3. HMA sponges follow a trade-off between lower pumping rates and better “cultivating” the microbiome. Differences in all mechanisms together are required to produce a good representation of the data. (A) Here, the difference between HMA and LMA is only that LMAs have 2.5 higher pumping rates. This results in reversed patterns, with HMA displaying lower abundance, richness, and evenness. (B) Then, on top of the differences in pump rates, considering resource allocation alone (100 times higher for HMA), the patterns are qualitatively well-represented, but the abundance ratio is lower and other differences are exaggerated. (C) Considering only selection on top of pumping rate (HMA selection capacity of 0.2 and LMA of 0.01), without differences in resource allocation, the patterns do not reflect observation of natural microbiomes. HMA richness becomes very low and total abundances are roughly on the same level. For this analysis, we simulated 6 LMA and 6 HMA species, each with 3 samples.

**Supplementary figure 3.**
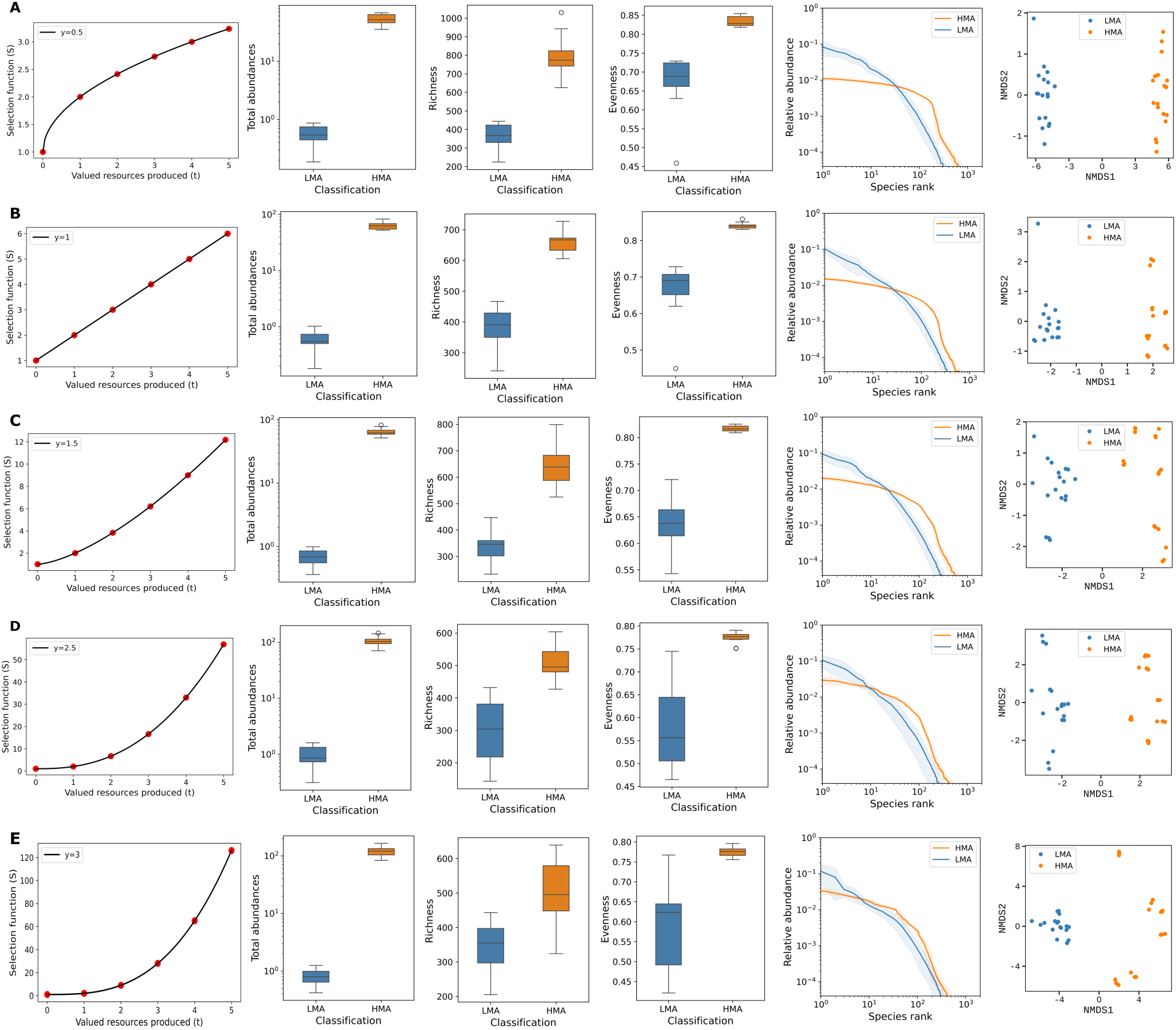
Sensitivity analysis on the selection function. Model parameters and plots are the same as presented in Fig 2 and Fig 3. Each row shows results for a particular curve, defined by the selection exponent *y* in Eq 3 with value equal to 2 in the main model. Values are (A) *y* = 0.5, (B) *y* = 1 (linear), (C) *y* = 1.5, (D) *y* = 2.5, and (E) *y* = 3. For this analysis, we simulated 6 LMA and 6 HMA species, each with 3 samples.

**Supplementary figure 4.**
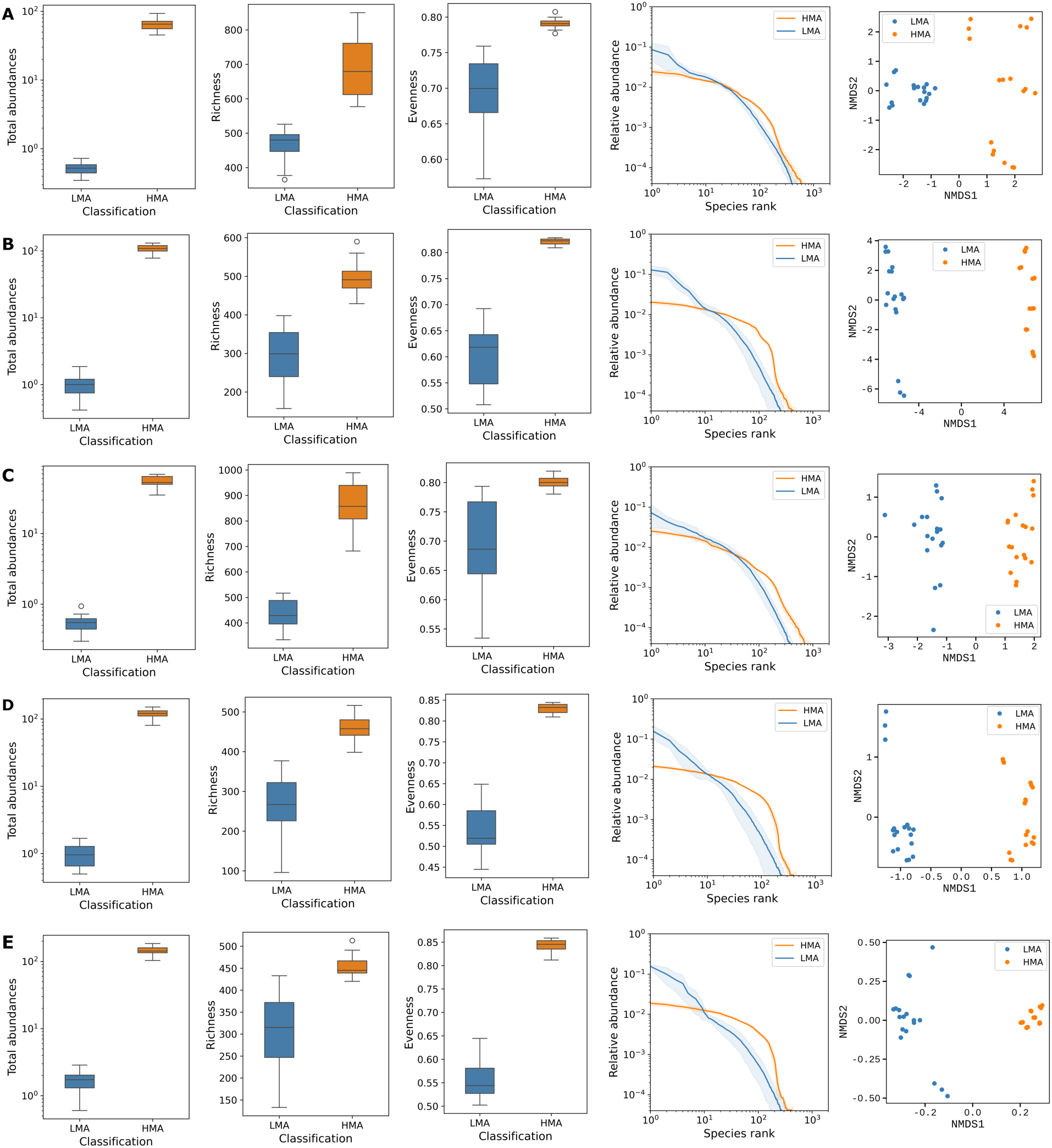
Sensitivity analysis on the resource values. Model parameters and plots are the same as presented in Fig 2 and Fig 3. Each row shows results for a particular configuration of fixed and random resources valued by hosts (as explained in Box 2). The main model uses (*q*_1_ = 15, *q*_2_ = 15) and we tested different proportions and values: (A) (*q*_1_ = 15, *q*_2_ = 0), (B) (*q*_1_ = 15, *q*_2_ = 30), (C) (*q*_1_ = 5, *q*_2_ = 15), (D) (*q*_1_ = 30, *q*_2_ = 15), and (E) (*q*_1_ = 30, *q*_2_ = 30). For this analysis, we simulated 6 LMA and 6 HMA species, each with 3 samples.

**Supplementary figure 5.**
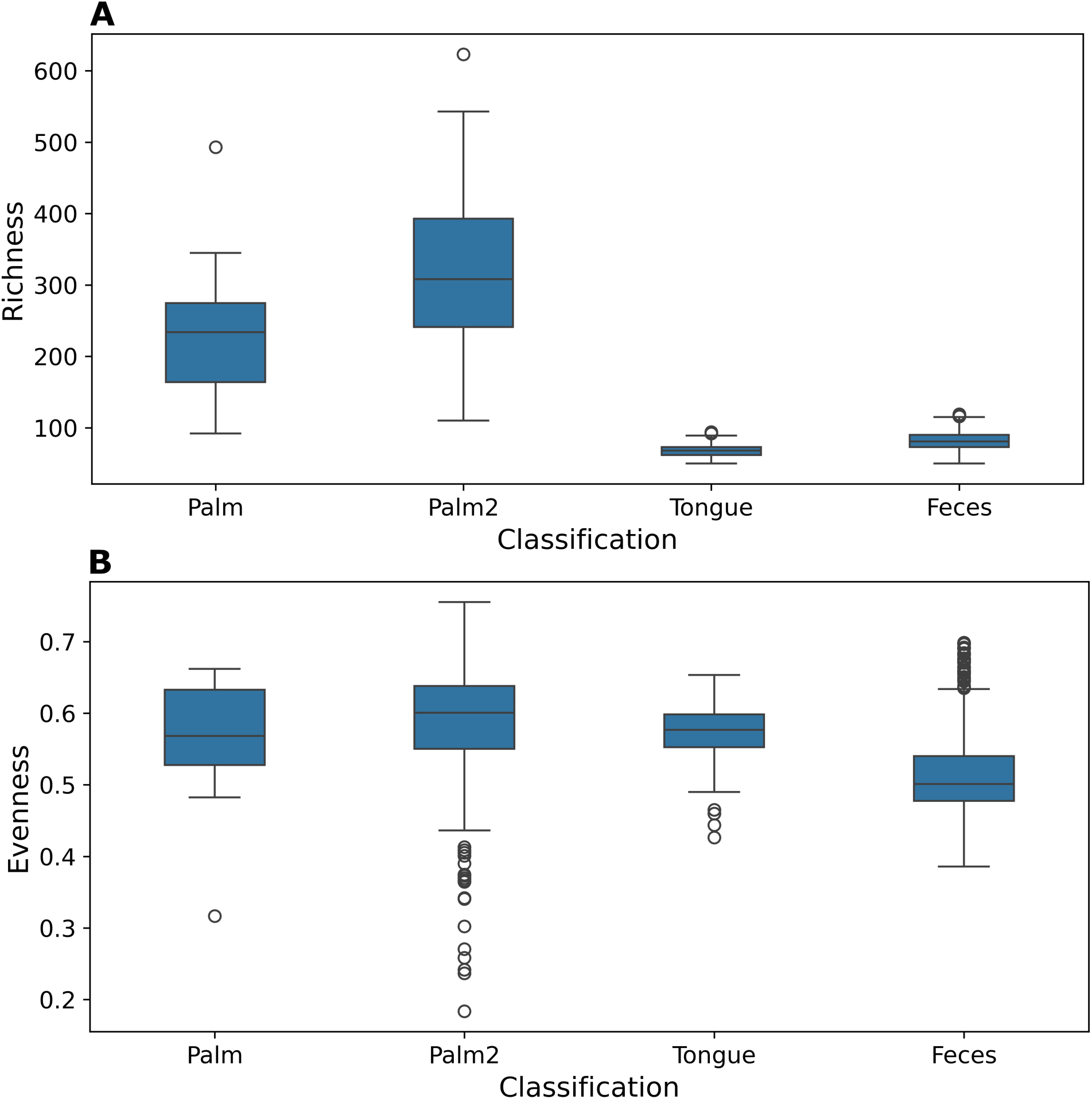
Detailed distributions for human microbiome data. The same analysis summarised in Fig 3 but shown for each microbiome separately. Groups belong to a particular microbiome of a single individual, with different samples taken in different moments (timeseries). (A) and (B) show respectively distributions of richness and evenness.

